# Prenatal methadone exposure selectively alters protein expression in primary motor cortex: implications for synaptic function

**DOI:** 10.1101/2022.12.21.521486

**Authors:** David L. Haggerty, Gregory G. Grecco, Jui-Yen Huang, Emma H. Doud, Amber L. Mosley, Hui-Chen Lu, Brady K. Atwood

## Abstract

As problematic opioid use has reached epidemic levels over the past two decades, the annual prevalence of opioid use disorder (OUD) in pregnant women has also increased 333%. Yet, how opioids affect the developing brain of offspring from mothers experiencing OUD remains understudied and not fully understood. Animal models of prenatal opioid exposure have discovered many deficits in the offspring of prenatal opioid exposed mothers, such as delays in the development of sensorimotor function and long-term locomotive hyperactivity. In attempt to further understand these deficits and link them with protein changes driven by prenatal opioid exposure, we used a mouse model of prenatal methadone exposure (PME) and preformed an unbiased multi-omic analysis across many sensoriomotor brain regions known to interact with opioid exposure. The effects of PME exposure on the primary motor cortex (M1), primary somatosensory cortex (S1), the dorsomedial striatum (DMS), and dorsolateral striatum (DLS) were assessed using quantitative proteomics and phosphoproteomics. PME drove many changes in protein and phosphopeptide abundance across all brain regions sampled. Gene and gene ontology enrichments were used to assess how protein and phosphopeptide changes in each brain region were altered. Our findings showed that M1 was uniquely affected by PME in comparison to other brain regions. PME uniquely drove changes in M1 glutamatergic synapses and synaptic function. Immunohistochemical analysis also identified anatomical differences in M1 for upregulating the density of glutamatergic and downregulating the density of GABAergic synapses due to PME. Lastly, comparisons between M1 and non-M1 multi-omics revealed conserved brain wide changes in phosphopeptides associated with synaptic activity and assembly, but only specific protein changes in synapse activity and assembly were represented in M1. Together, our studies show that lasting changes in synaptic function driven by PME are largely represented by protein and anatomical changes in M1, which may serve as a starting point for future experimental and translational interventions that aim to reverse the adverse effects of PME on offspring.

## Introduction

As problematic opioid use has reached epidemic levels over the past two decades, the annual prevalence of opioid use disorder (OUD) in pregnant women has also increased 333% (Haight, Ko et al. 2018). A product of this increased opioid use has also lead to a seven-fold increase in neonates experiencing opioid withdrawal syndrome (NOWS) following prenatal opioid exposure (POE) (Epstein, Bobo et al. 2013, Patrick, Davis et al. 2015, Ko, Patrick et al. 2016, Ko, Wolicki et al. 2017, Duffy, Wright et al. 2018, Winkelman, Villapiano et al. 2018, Honein, Boyle et al. 2019). POE negatively impacts neurodevelopment, producing dysfunction of motor control and persistent behavioral hyperactivity, with potential long-term negative effects on quality of life (Sundelin Wahlsten and Sarman 2013, Lee, Woodward et al. 2019, Yeoh, Eastwood et al. 2019). Recent data also show that POE neonates who display NOWS symptoms at birth have better outcomes than POE neonates that did not display NOWS, likely a function of the treatments used to alleviate NOWS potentially having lasting protective effects (Leyenaar, Schaefer et al. 2021). Despite the proven potential for treatments to improve the lives of neonates with POE, there has been a lack of new treatments as current first-line treatments for both pregnant women with OUD and neonates displaying NOWS remain opioid maintenance therapies, such as methadone and buprenorphine (2017, Ghazanfarpour, Najafi et al. 2019, Tang, Ng et al. 2021, Zankl, Martin et al. 2021). As a consequence, it is difficult to separate the pharmacology of POE and NOWS treatments as well as other environmental factors, both independently and in combination with each other across the human lifespan. This complexity raises questions regarding the specific impacts of opioids and NOWS treatments alone. Ultimately, it is unknown exactly how POE impacts the brain and behavioral development of these neonates as they mature (Brogly, Saia et al. 2018, Rose-Jacobs, Trevino-Talbot et al. 2019, Labella, Eiden et al. 2021).

Previous rodent studies have revealed some important findings regarding the specific impacts of POE on neurobiological function and/or behavioral development through the lifespan (Grecco and Atwood 2020, Boggess and Risher 2022). However, many of these prior studies have inadequately modelled human clinical cases, making their findings difficult to interpret and hindering translation to human therapeutics. Recently, our lab has developed a translational mouse model that more closely resembles the typical pattern of opioid use in a pregnant woman who is first dependent on oxycodone, then begins methadone maintenance treatment, and subsequently becomes pregnant while maintained on methadone (Grecco, Mork et al. 2021). This prenatal methadone exposure (PME) model produces many outcomes reminiscent of human neonatal opioid withdrawal. In this model, we found that PME reduces physical growth in offspring persisting into adolescence and delays the development of sensorimotor function during the pre-weaning period. Furthermore, PME produced locomotor hyperactivity that persisted into adolescence as well.

The potential neural mechanisms that mediate the sensorimotor deficits and hyperactivity observed in our model are diverse. We previously characterized PME-induced physiological and biochemical changes in the dorsal striatum, a brain region that integrates input from sensorimotor cortices (Grecco, Munoz et al. 2022). We have also described the effects of PME on the biochemical, anatomical and electrophysiological properties of the primary somatosensory cortex (S1) (Grecco, Huang et al. 2022). The effects we measured in these brain regions likely contribute to the sensorimotor behavioral changes we observed as a result of PME. However, the primary motor cortex (M1) likely also contributes, given its central role in the preparation, execution, and adaptation of motor movements (Peters, Liu et al. 2017, Svoboda and Li 2018). Indeed, we showed that PME alters the physiological parameters of layer 5 pyramidal neurons within M1 with similar effects in both males and females (Grecco, Mork et al. 2021). Specifically, PME enhanced excitatory glutamatergic input to these neurons from outer M1 layers and enhanced voltage sag, an electrophysiological property related to the function of hyperpolarization-activated cyclic nucleotide-gated (HCN) channels (Sheets, Suter et al. 2011). However, in our previous study of M1 we failed to explore the biochemical changes produced by PME in M1 that could underlie the physiological changes it induced. Thus, we sought to profile the protein expression and phosphorylated protein states in this brain region and compare these changes to those in S1 and the lateral (DLS) and medial (DMS) divisions of the dorsal striatum. By tracking the protein expression and protein phosphorylation states altered by PME across distinct brain regions, we can learn if PME has differential effects across different brain regions, and further elucidate how POE uniquely alters infants in hopes of refining our approaches to treating NOWS and mitigate POE-induced long-lasting negative outcomes.

## Materials and Methods

### Animals

Animal care and research were conducted in accordance with guidelines established by the National Institutes of Health and protocols were approved by the Indiana University School of Medicine Institutional Animal Care and Use Committee. Eight-week-old female C57BL/6J mice were acquired from Jackson Laboratories (Bar Harbor, Maine), single housed, and randomly assigned to either saline (10 mL/kg) or oxycodone treatments. Oxycodone dependence was induced by a dose-ramping procedure with a dose of 10 mg/kg administered on pregestational day (PG) 14, 20 mg/kg on PG13, and then maintained on 30 mg/kg for PG12-6. All saline or oxycodone doses were administered subcutaneously twice daily at least 7 hr apart. On PG5, oxycodone-treated mice were transitioned to 10 mg/kg methadone while saline-treated animals continued to receive saline injections (s.c. b.i.d.). All oxycodone and methadone solutions were prepared in saline. Oxycodone and methadone were obtained from the National Institute on Drug Abuse Drug Supply Program. Five days following initiation of methadone treatment, an 8-week-old C57BL/6J male mouse (also acquired from Jackson Laboratories) was placed into the cage of each female for four days. Mucous plugs were assessed each morning to approximate gestational day (G) 0.

Cages were examined for the presence of pups at the time of each morning and afternoon injection, and the day of birth was designated postnatal day (P) 0. Only litters between three and eight pups were used in subsequent studies of offspring. To maintain a sufficient level of methadone exposure to the newborn pups during P0-P7, the dams were maintained at their highest gestational dose after giving birth and up to P7. After P7, the dose of methadone administered to dams was adjusted to their body weight. All treatments to dams were discontinued at weaning (∼P28). For more detailed methods on methadone exposure and the associated behavioral changes, please see our previous publication that defines the PME model (Grecco, Mork et al. 2021).

Early adolescent offspring (P21–P36) were used for the proteomics and immunohistochemical studies. Sample preparation, mass spectrometry analysis, bioinformatics, and data evaluation for quantitative proteomics and phosphoproteomics experiments were performed in collaboration with the Indiana University Proteomics Core similar to our previous studies (Grecco, Haggerty et al. 2021, Grecco, Huang et al. 2022). Our previous studies of motor behavior and M1 physiology did not reveal sex differences. In addition, statistical power calculations indicated that the number of samples available to us for our proteomics assessments would not be sufficient to detect sex x prenatal exposure interactions. Therefore, in this study we limited our comparisons to prenatal treatment alone but included data from both males and females in each dataset. We have illustrated data that came from males and females throughout the paper so that the reader might discern whether future, more powered studies may reveal sex differences.

### M1 Proteomics and Phosphoproteomics

#### Protein Preparation

For proteomics and phosphoproteomics, we followed similar methods on which we have previously published, with the exception that we used the more recently updated mouse reference proteome database (Grecco, Haggerty et al. 2021, Grecco, Huang et al. 2022). Animals were rapidly decapitated without anesthesia by blinded researcher and tissue was dissected. Slices were cut in a 0.5 mm coronal mouse brain matrix and whole M1, S1, DMS, and DLS were dissected from each slice. Tissue was immediately snap frozen in isopentane on dry ice and stored at −80°C. Flash frozen brain lysates were homogenized using a BeadBug™ 6 (Benchmark scientific Cat No: D1036, 3 mm zirconium beads Cat No: D1032-30, 10 rounds of 30 × 30 s,4°C) in 1 mL of 8 M urea (CHEBI: 16199) in 100 mM Tris, pH 8.5 (CHEBI: 9754). Samples were then sonicated on a Bioruptor® sonication system (Diagenode Inc. United States, North America Cat No: B01020001) with 30 s/30 s on/off cycles for 15 min in a water bath at 4°C. After subsequent centrifugation at 14,000 rcf for 20 min, protein concentrations were determined by Bradford protein assay (BioRad Cat No: 5000006). 100 µg equivalent of protein from each sample were then treated with 5 mM tris (2-carboxyethyl)phosphine hydrochloride (Sigma-Aldrich Cat No: C4706) to reduce disulfide bonds and the resulting free cysteine thiols were alkylated with 10 mM chloroacetamide (Sigma Aldrich Cat No: C0267). Samples were diluted with 50 mM Tris pH 8.5 (Sigma-Aldrich Cat No: 10812846001) to a final urea concentration of 2 M for overnight Trypsin/Lys-C digestion at 35°C (1:100 protease:substrate ratio, Mass Spectrometry grade, Promega Corporation, Cat No: V5072 (Levasseur, Yamada et al. 2019, Li, Van Vranken et al. 2020).

#### Peptide Purification and Labeling

We performed peptide purification and labeling according to previously published methods (Grecco, Haggerty et al. 2021, Grecco, Huang et al. 2022). To summarize our previous methods, digestion was halted by addition of 0.3% final v/v trifluoroacetic acid (TFA), and peptides were desalted on Waters Sep-Pak® Vac cartridges (Waters™ Cat No: WAT054955) with a wash of 1 mL 0.1% TFA followed by elution in 0.6 mL of 70% acetonitrile 0.1% formic acid (FA). Peptides were dried by speed vacuum and resuspended 50 mM triethylammonium bicarbonate. Peptide concentrations were checked by Pierce Quantitative colorimetric assay (Cat No: 23275). The same amount of peptide from each sample was then labeled for 2 hours at room temperature, with 0.5 mg of Tandem Mass Tag Pro (TMTpro) reagent (16-plex kit, manufactures instructions Thermo Fisher Scientific, TMTpro™ Isobaric Label Reagent Set; Cat No: 44520, lot no. VI310352) (Li, Van Vranken et al. 2020). Labeling reactions were quenched with 0.3% hydroxylamine (final v/v) at room temperature for 15 min. Labeled peptides were then mixed and dried by speed vacuum. The TMT-labeled peptide mix was desalted to remove excess label using a 100 mg Waters SepPak cartridge, eluted in 70% acetonitrile, 0.1% formic acid and lyophilized to dryness.

#### Phosphopeptide Enrichment

Phosphopeptides were enriched from the mixed, labeled peptides on one spin tip from a High-Select™ TiO2 Phosphopeptide Enrichment Kit (capacity of 1–3 mg; Thermo Fisher Scientific, catalog A32993). After preparing spin tips, labeled and mixed peptides were repeatedly applied to the TiO2 spin tip, eluted, and immediately dried as per manufacturer’s instructions. Prior to LC/MS/MS the phosphopeptides were resuspended in 25 µL 0.1% formic acid. The flow through from each tip was saved for global proteomics.

#### High pH Basic Fractionation

The phosphopeptide enrichment step flow through and washes were dried down and approximately 120 µg peptides were fractionated using the TMT fractionation protocol of Pierce high pH basic reversed-phase peptide fractionation kit (Thermo Fisher Scientific™ Cat no 84858; with a wash of 5% acetonitrile, 0.1% triethylamine (TEA) followed by elution in 10%, 12.5%, 15%, 17.5%, 20%, 22.5%, 25%, 30%, and 70% acetonitrile, all with 0.1% TEA). Fractions were dried down and resuspended in 50 µL 0.1% FA prior to online LC-MS.

#### Nano-LC-MS/MS Analysis

We performed nano-LC-MS/MS analysis with previously published methods (Grecco, Haggerty et al. 2021, Grecco, Huang et al. 2022). To summarize our previous methods, nano-LC-MS/MS analyses were performed on an EASY-nLC™ HPLC system (SCR: 014993, Thermo Fisher Scientific) coupled to Orbitrap Fusion™ Lumos™ mass spectrometer (Thermo Fisher Scientific). One fifth of the phosphopeptides and one tenth of each global peptide fraction was loaded onto a reversed phase EasySpray™ C18 column (2 μm, 100 Å, 75 μm × 25 cm, Thermo Scientific Cat No: ES902A) at 400 nL/min. Peptides were eluted from 4 to 28% with mobile phase B [Mobile phases A: 0.1% FA, water; B: 0.1% FA, 80% Acetonitrile (Fisher Scientific Cat No: LS122500)] over 160 min; 28%–35% B over 5 min; 35–50% B for 14 min; and dropping from 50 to 10% B over the final 1 min. The mass spectrometer method was operated in positive ion mode with a 4 s cycle time data-dependent acquisition method with advanced peak determination and Easy-IC (internal calibrant). Precursor scans (m/z 400-1750) were done with an orbitrap resolution of 120,000, RF lens% 30, maximum inject time 50 ms, standard AGC target, including charges of 2–6 for fragmentation with 60 s dynamic exclusion. MS2 scans were performed with a fixed first mass of 100 m/z, 34% fixed CE, 50000 resolution, 200% normalized AGC target and dynamic maximum IT. The data were recorded using Thermo Fisher Scientific Xcalibur (4.3) software (Thermo Fisher Scientific Inc.)

#### Proteome and Phosphoproteome Data Processing

We performed M1, S1, DLS, and DMS data processing similar to our previously published methods with the exception that we analyzed our resulting M1 and previous RAW files in Proteome Discover™ 2.5 (Thermo Fisher Scientific, RRID: SCR_014477) with a more recent *Mus musculus* UniProt FASTA (both reviewed and unreviewed sequences, downloaded 05/13/2022, 84,754 sequences plus common contaminants (71 sequences)) (Grecco, Haggerty et al. 2021, Grecco, Huang et al. 2022). SEQUEST HT searches were conducted with a maximum number of 3 missed cleavages; precursor mass tolerance of 10 ppm; and a fragment mass tolerance of 0.02 Da. Static modifications used for the search were, 1) carbamidomethylation on cysteine (C) residues; 2) TMTpro label on lysine (K) residues and the N-termini of peptides. Dynamic modifications used for the search were TMTpro label on N-termini of peptides, oxidation of methionines, phosphorylation on serine, threonine or tyrosine, and acetylation, methionine loss or acetylation with methionine loss on protein N-termini. Percolator False Discovery Rate was set to a strict setting of 0.01 and a relaxed setting of 0.05. IMP-ptm-RS node was used for all modification site localization scores. Values from both unique and razor peptides were used for quantification. In the consensus workflows, peptides were normalized by total peptide amount with no scaling. Quantification methods utilized TMTpro isotopic impurity levels available from Thermo Fisher Scientific. Reporter ion quantification was allowed with S/N threshold of 7 and co-isolation threshold of 50%. Data shown is for PME/PSE abundance value ratios (AR). Resulting grouped abundance values for each sample type, AR values; and respective p-values (t-test) from Proteome Discover™ were exported to Microsoft Excel. Full datasets are provided in the Supplementary Material. The raw data files can be found at https://github.com/dlhagger/PME-MultiRegion-Proteomics.

### Immunohistochemistry

We performed immunohistochemistry and its analysis similar to previously published methods (Grecco, Haggerty et al. 2021, Grecco, Huang et al. 2022). To summarize our previous methods, animals were anesthetized with isoflurane and perfused with 4% paraformaldehyde prepared in PBS for 10 mins at a pump rate of ∼2 mL/min. Fixed brains were sectioned into 100 μm sections in the coronal plane (between bregma −0.1 and −1.94 mm) using a Leica VT-1000 vibrating microtome (Leica Microsystems) and stored in antigen preserved solution (PBS, 50% ethylene glycol and 1% polyvinyl pyrrolidone) at −20°C. For synaptic marker (VGAT, Gephyrin, VGluT1, and PSD95) staining, sections were permeabilized with 2% Triton X100, then incubated with a blocking solution (3% normal goat serum prepared in PBS with 0.3% Triton X-100) and then incubated overnight with primary antibody prepared in blocking solution (See Table 1 for concentration and source). An appropriate secondary antibody conjugated with an Alexa series fluorophore was used to detect the primary antibody. DAPI (100 ng/ml, Thermo Fisher) was included in the secondary antibody solution to stain nuclei.

**Table 1.** Antibody Descriptions.

| Antibodies | Host | Source | RRID# | Titer |
| --- | --- | --- | --- | --- |
| VGAT | Rabbit | SYSY (131-002) | AB_887871 | 1:2000 |
| Gephyrin | Mouse | SYSY(147-021) | AB_2232546 | 1:1000 |
| PSD95 | Rabbit | Thermo Fisher (51-6900) | AB_2533914 | 1:2000 |
| VGluT1 | Guinea Pig | Millipore (AB5905) | AB_2301751 | 1:2000 |

For imaging synaptic marker staining we followed similar methods which we have previously published (Grecco, Huang et al. 2022). Z-stack confocal images were acquired from both hemispheres with a Nikon A1 confocal microscope with a 60X/NA1.4 objective at 3 times software zoom or Leica SP8 confocal microscope with a 63X/NA1.2 objective at 2.5 times software zoom. The Z-stacks were taken at 0.1 µm intervals (for VGAT + Gephyrin) or 0.2 µm (for VGluT1 + PSD95), and 2–4 µm-total thickness was imaged. Two images from each hemisphere and both hemispheres were imaged per animal. We utilized Imaris (Bitplane, Zurich, Switzerland) to quantify synaptic punctate at the three-dimensional level and establish the data analysis workflow to quantify the synaptic number according to the published literature (Fogarty, Hammond et al. 2013, Kuljis, Park et al. 2019, Simhal, Zuo et al. 2019, Zhou, Chen et al. 2019). The volume occupied by nuclei and vasculature varied within each image, thus robustly impacting the density of synaptic marker quantification. To accurately estimate the neuropil occupied volume, we first used surface module to create the surface objects of nuclei and vasculature-like structure. Next, the gephyrin- or PSD95-channel was further masked by nuclei and vasculature objects to exclude the volume occupied by nuclei and vasculature. The post-masked gephyrin or PSD95 channel was used to generate a surface object containing the volume (neuropil object) to be analyzed. For spot detection, we followed similar procedures and parameter settings as described before (Fogarty, Hammond et al. 2013, Kuljis, Park et al. 2019, Simhal, Zuo et al. 2019, Zhou, Chen et al. 2019). Specifically, the pre-synaptic (VGluT1 and VGAT) and postsynaptic (gephyrin and PSD95) punctate were detected by Imaris spot module with 0.5 µm and 0.3 µm diameter according to the published literature. In general, the diameter for synaptic puncta is between 0.25–0.8 µm (Kim and Sheng 2009, Dumitriu, Berger et al. 2012). To find the optimal detecting threshold for spot detection, we first manually defined the detecting threshold for one image from each animal. The threshold that detected most synaptic punctates without creating artifacts was applied to analyze all images and generated the spot layer for each synaptic marker. Only synaptic punctates inside the neuropil-object were used for subsequence analysis. Next, we determine how many pre-synaptic spots were directly opposed to postsynaptic spots (defined as synapse at anatomical level) with the distance 0.5 µm. The juxtaposed synaptic punctate of VGluT1/PSD95 and VGAT/gephyrin were defined as intracortical excitatory and inhibitory neurochemical inputs. Synaptic density was calculated as the number of synapses detected in a dataset over the volume of the dataset. All image acquisition and data analysis were performed in a blinded manner.

### Gene Ontology Enrichment Analysis

All analyses are presented as PME relative to PSE (log2 abundance ratios of PME/PSE). For protein expression dendrograms, all differentially abundant (p < 0.05) proteins and phosphopeptides were normalized across animal by their associated UniProt Accession numbers and then hierarchically clustered using the Voorhees algorithm. Predicted protein-protein associative networks and functional enrichments were computed by loading the associated GeneIDs and log2 abundance ratios of differentially abundant proteins and phosphopeptides into STRING (Szklarczyk, Gable et al. 2021). Protein-protein and phosphopeptide-phosphopeptide interactions were visualized using the highest confidence interaction score (0.9) with disconnected nodes hidden from the visualization. The interaction network was then kmeans clustered (n=5) and assigned a pseudo-color to represent each subcluster that was post-hoc defined by the top 1 or 2 strongest KEGG pathway terms returned by each subcluster’s identity. For overrepresentation analysis of Gene Ontology (GO), the STRING analysis output was filtered to return only significant (p < 0.05) terms for the following classifications: GO Process, GO Function, KEGG, and Reactome. The similarity of GeneIDs and Gene Ontology terms for M1 compared to all other brain regions were computed using cosine similarity between the lists of GeneIDs or GO terms to demonstrate how closely related these identifiers were between brain regions. The full results of the GO analysis are provided in the Supplementary Material and at https://github.com/dlhagger/PME-MultiRegion-Proteomics.

### Statistics and Data Presentation

Data are graphically presented as dot plots displaying all individual data points with the inclusion of box plots that display interquartile ranges as whiskers. Individual data points that fall beyond the interquartile range are noted with a diamond next to the individual data point. The level of significance was a priori set at p < 0.05. All experiments were performed using both male and female offspring. To minimize potential litter effects in all completed studies, no more than two males and females per litter were utilized for any study. Immunostaining studies were not sufficiently powered to detect sex differences with sex considered as a factor (determined by power analysis), and therefore the data presented here are collapsed on sex as a factor. Immunostaining statistical analyses were conducted using pingouin (Vallat 2018). T-tests, with Welch’s correction where appropriate, were used for analyzing all immunostaining data.

## Results

### Differential Protein and Phosphopeptide Expression

To determine the possible unique alterations that underlie dysfunctional development in PME offspring that occur specifically in the motor cortex, we collected brain punches of M1 cortices from adolescent male and female PME and PSE offspring for quantitative proteomic and phosphoproteomic analyses. In M1 alone, we identified 5,974 proteins and 5,794 phosphopeptides in offspring that were altered by PME. For M1, 1,248 proteins and 391 phosphopeptides were differentially abundant (**Figure 1A and B**). For the M1 proteome, a substantial downregulation of proteins was observed, with 200 proteins being up-regulated and 1,048 proteins being down-regulated. Whereas for the M1 phosphoproteome, 199 phosphopeptides were up-regulated, and 192 phosphopeptides were down-regulated. Clustermaps of the differentially abundant proteins show that PME and PSE animals are most dissimilar, yet male and female animals are most dissimilar for differentially abundant phosphopeptides (**Figure 1C and D**). For a full detailed list of differentially abundant proteins, phosphopeptides, and clustermaps by individual sample, please see **Supplemental Figure 1** and the Supplementary Files for M1 regions. Together, these data suggest that PME alters more global protein abundances than phosphopeptides within the motor cortex.

**Figure 1:**
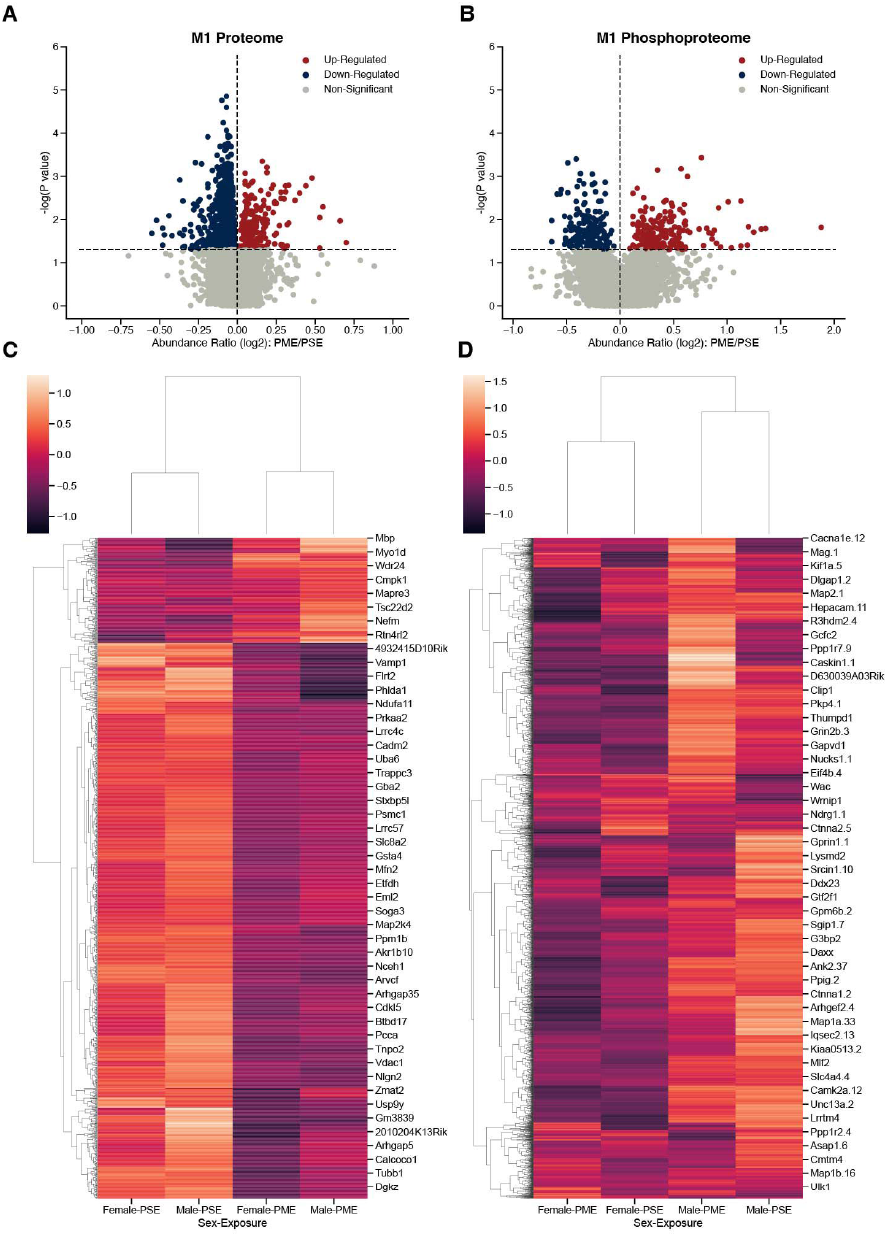
Differential protein and phosphopeptide expression in the primary motor cortex (M1) of prenatal methadone exposed offspring. Volcano plots for the differential proteome (A) and phosphoproteome (B) with blue circles representing individual proteins/phosphopeptides decreased in PME vs. PSE and red circles representing individual proteins/phosphopeptides increased in PME vs. PSE which reach the level of significance. Clusterplots of differentially abundant proteins (C) and phosphopeptides (D) with example GeneIDs. (n = 8 (4M:4F) PME, 8 PSE (4M:4F)).

### Protein-Protein and Phosphopeptide-Phosphopeptide Interaction Networks

To further understand the differences in the proteome and phosphoproteome of the primary motor cortex, differentially abundant proteins and phosphopeptide interaction networks were computed. By using the highest confidence interactions and showing only connected nodes, the protein interaction network was constructed then k-means clustered (k = 5) to uncover the identifies of subclusters within the network (**Figure 2A**). The many M1 proteomic interactions show major KEGG defined clusters for alterations in autophagy, MAPK/Rap1/Ras signaling, metabolic pathways alterations, and changes in glutamatergic signaling at synapses. For M1 phosphoproteomics, a substantially smaller interaction network did not return significant KEGG defined subclusters, therefore a smaller k-means clustering (k=3) of Reactome terms was used to uncover the identities of network subclusters (**Figure 2B**). Altered mRNA splicing, membrane trafficking, and presynaptic depolarization and calcium channel opening were identified as the strongest clusters for the M1 phosphoproteome. Interestingly, both networks contain strong alterations in synapses, as well as calcium channel alterations associated with neuronal activity.

**Figure 2:**
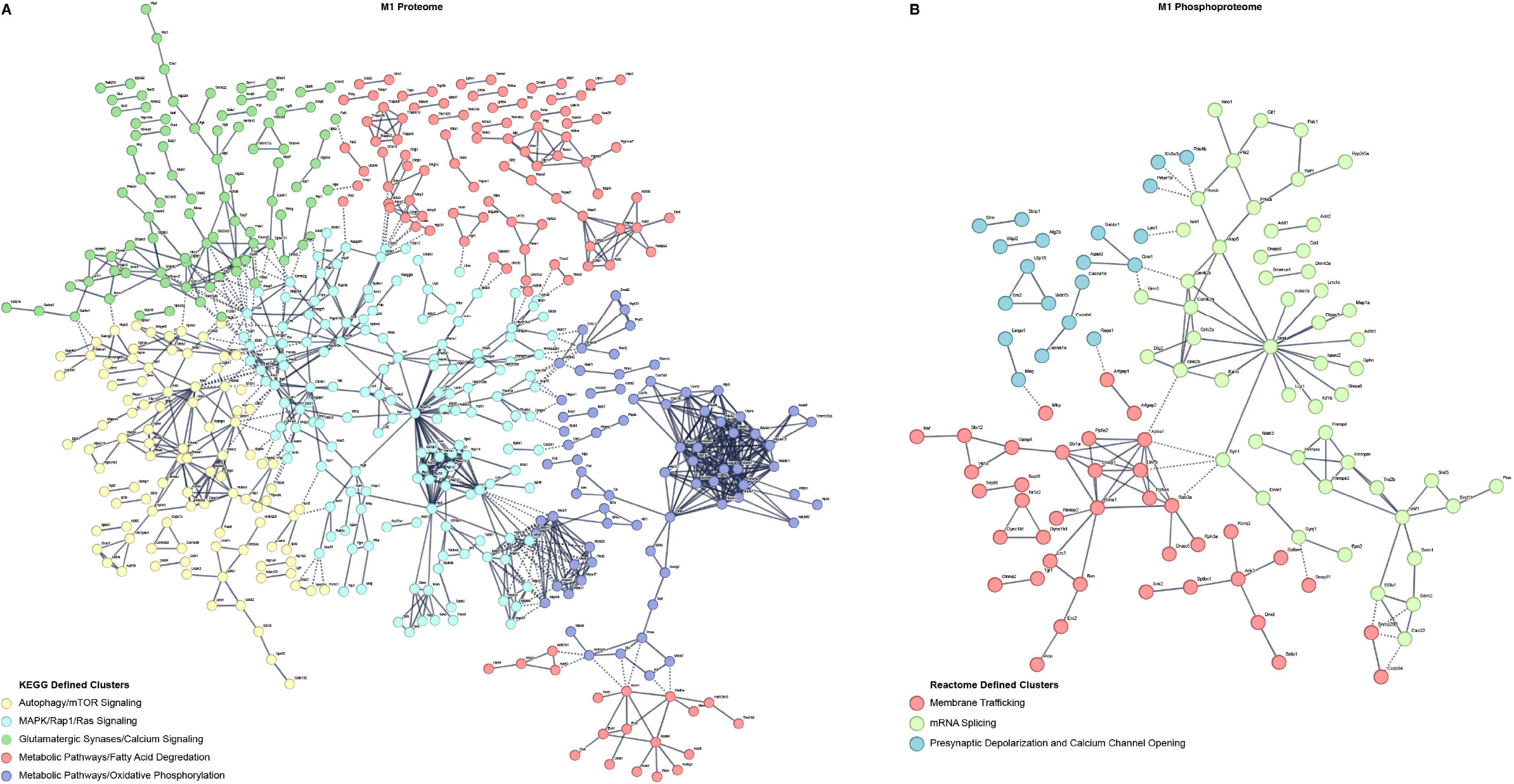
Differential protein and phosphopeptide interaction networks in the primary motor cortex (M1) of prenatal methadone exposed offspring. Interaction networks for the differential proteome (A) and phosphoproteome (B). Interaction networks were k-means clustered and the strongest KEGG or Reactome term of the subcluster was used to define the individual subcluster.

### Gene and Gene Ontology Functional Enrichments

Although the number of individual proteins and the abundance change magnitudes were larger in the global proteome than the enriched phosphoproteome PME in M1, we sought to further investigate the overlap between PME-induced changes in the proteome and phosphoproteome. By looking at the 89 overlapping associated GeneIDs from differentially abundant proteins and phosphopeptides and returning the 5 greatest positive and negative changes in log_2_(abundance ratios), there exists an overlap in targets such as Grin2a, Dlgap3, Ank2, Nrgn, and Mobp (**Figure 3A**). These genes are all known to regulate synaptic organization, activity, and are known targets in disorders associated with aberrant neuronal activity or synapse loss (Bulat, Rast et al. 2014, Wan, Ade et al. 2014, Hegyi 2017, Rasmussen, Rasmussen et al. 2017, Amador, Bostick et al. 2020, Pereira, Janelidze et al. 2021).

**Figure 3:**
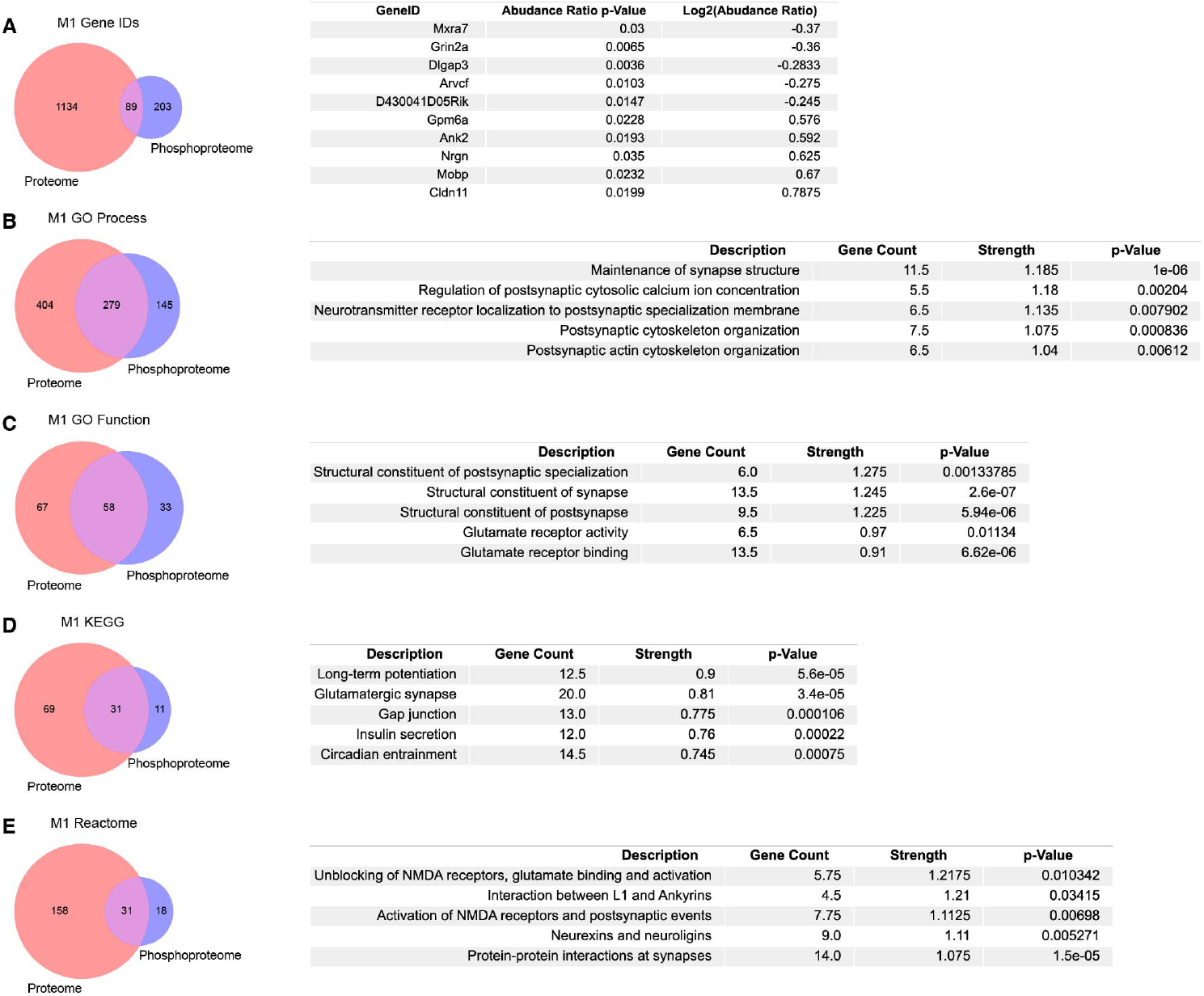
Overlap in the proteome, phosphoproteome, and gene ontology enrichments between prenatal methadone exposed primary motor cortex (M1). (A) Of 1426 differentially abundant proteins and phosphopeptides matched to their corresponding GeneIDs, 89 were both similarly altered in the proteome and phosphoproteome with the top 5 largest positive and negative log_2_(abundance ratio) listed. Overlap between M1 proteins and phosphopeptides for gene ontology enrichments for GO Process (B), GO Function (C), KEGG (D), and Reactome (E), listed with the top 5 terms for each enrichment, sorted by the strength of enrichment.

We similarly analyzed overlap in PME-altered expression of M1 proteins and phosphopeptides using gene ontology databases to further characterize the functional changes beyond specific associated gene targets using STRING (Szklarczyk, Gable et al. 2021). The overlap between GO Processes showed 279 matching terms (**Figure 3B**). When ranking these GO Processes by their greatest strength, the 5 greatest terms all are associated with synapse maintenance, activity, and structure. Similarly, GO Function overlap in M1 shows 58 terms that are shared with the top 5 strongest terms all relating to structural changes at synapses and glutamate receptor activity (**Figure 3C**). 31 overlapping KEGG terms show that 3 of the top 5 strongest terms are enriched for synaptic plasticity and synaptic events, (**Figure 4D**) and 31 overlapping Reactome terms show that all of the strongest terms relate to synaptic changes, especially at glutamatergic synapses with NMDA receptors (**Figure 3E**). Together, the overlap between gene and gene ontology enrichments between protein and phosphopeptides overwhelmingly point to PME induced alterations in synaptic assemblies and activity, especially related to glutamatergic synapses.

**Figure 4:**
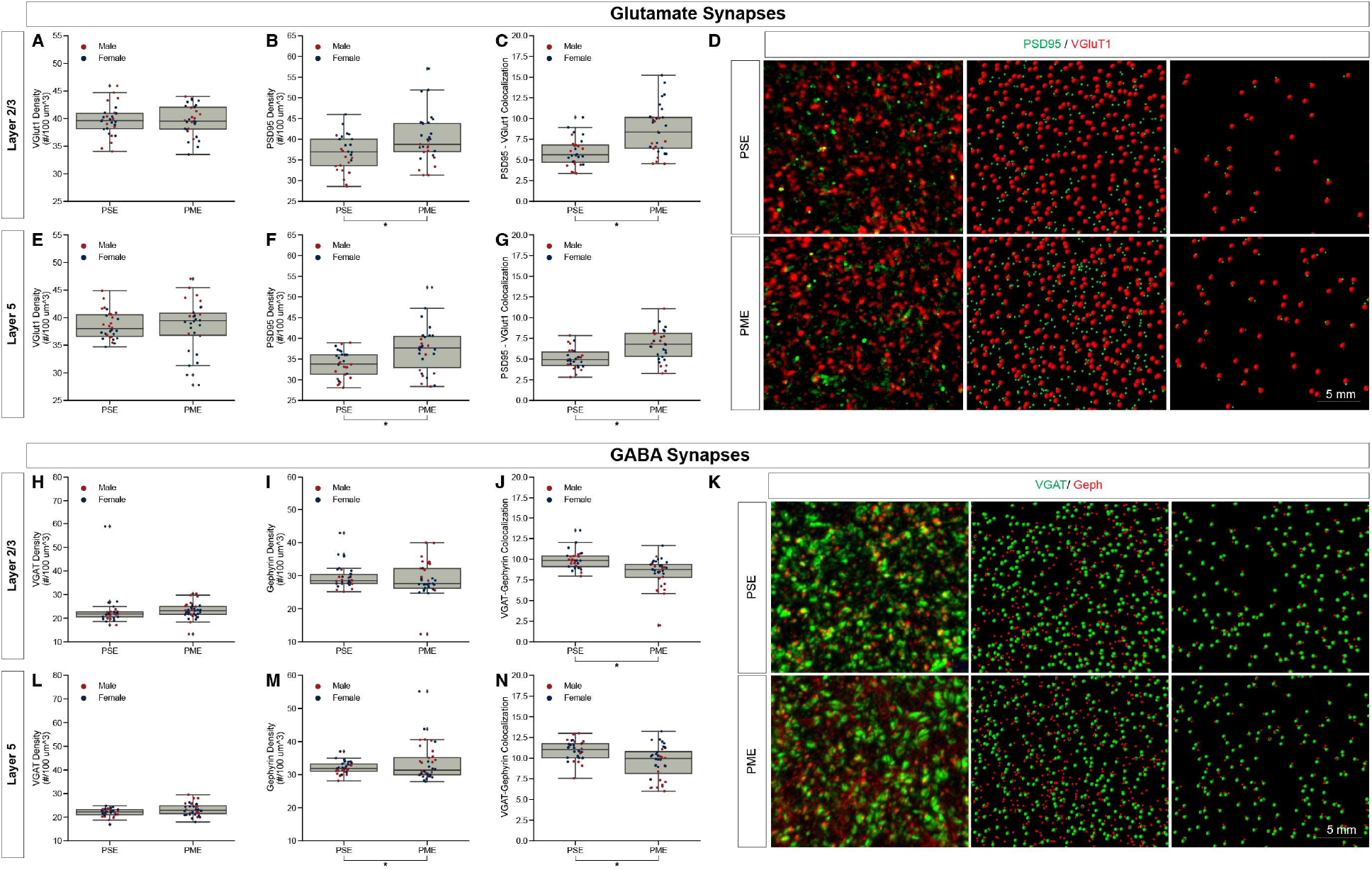
Neurochemical assessment of functional Glutamatergic and GABAergic synapses. In Layer 2/3 (A) vesicular glutamate transporter (VGlut1) density was unaltered, but PME increases (B) post-synaptic density (PSD95) density (t test, *p* = 0.0141) and (C) increases PSD95-VGlut1 colocalization (t test, *p* = 0.0001). An exemplary (D) confocal stack of PSD95 and VGlut1 double-stained image between PME and PSE. In Layer 5 (E) VGlut1 density was unaltered, but PME increases (F) PSD95 density (t test, *p* = 0.0031) and (G) increases PSD95-VGlut1 colocalization (t test, *p* = 0.0001). n = 8 (3M:5F) PME, 8 PSE (4M:4F). In Layer 2/3 both (H) vesicular GABA transporter (VGAT) density and (I) post-synaptic density (gephyrin) density was unaltered, but PME (J) decreases Gephyrin-VGAT colocalization (t test, *p* = 0.0005). An exemplary (K) confocal stack of gephyrin and VGAT double-stained image between PME and PSE. In Layer 5 (L) VGAT density was unaltered, but PME increases (M) Gephyrin density (t test, *p* = 0.0133) and (N) decreases gephyrin-VGAT colocalization (t test, *p* = 0.0005). n = 9 (4M:5F) PME, 8 PSE (4M:4F).

### Assessment of Glutamatergic Synaptic Markers

Given the large number of differentially abundant proteins and phosphopeptides associated with synaptic functioning and the enrichment in associated GeneIDs and terms associated with the synapse, we investigated both glutamatergic and GABAergic synapse density in layer 2/3 (L2/3) and layer 5 (L5) of M1 using the co-localization of presynaptic vesicular glutamate transporter (VGlut1) and postsynaptic protein PSD95 and co-localization of presynaptic vesicular GABA transporter (VGAT) and postsynaptic protein gephyrin, respectively. The co-localization of these makers was used to infer the presence of a “functional” glutamate or GABA synapse from anatomical data.

For glutamate synapses in L2/3 of M1, the density of VGlut1, a protein conventionally considered as marker of intracortical inputs, was unchanged by PME (**Figure 4A**). Yet, both PSD95 density (t test, *t*_(55)_ = 2.5351, *p* = 0.0141) and PSD95-VGlut1 co-localization (t test, *t*_(49)_ = 4.3503, *p* = 0.0001) were increased due to PME exposure (**Figure 4B,C**). A representative confocal stack of PSD95 and VGlut1 double-stained image between PME and PSE visualizes these differences (**Figure 4D**). Similarly, for glutamate synapses in L5 of M1, the density of VGlut1 was unchanged by PME (**Figure 4E**). Both, PSD95 density (t test, *t*_(58)_ = 3.0883, *p* = 0.0031) and PSD95-VGlut1 co-localization (t test, *t*_(58)_ = 4.0594, *p* = 0.0001) were also increased due to PME exposure (**Figure 4F,G**). For both L2/3 and L5, PME substantially increases the density of functional glutamatergic synapses in M1.

For GABA synapses in L2/3 of M1, the density of VGAT and gephyrin were both unchanged by PME (**Figure 4H,I**). Yet, VGAT-gephyrin co-localization (t test, *t*_(58)_ = −3.7058, *p* = 0.0005) was decreased due to PME exposure (**Figure 4J**). A representative confocal stack of gephyrin and VGAT double-stained image between PME and PSE visualizes these changes (**Figure 4K**). In L5, VGAT was also unaltered (**Figure 4L**). Gephyrin density was increased (t test, *t*_(60)_ = 2.5499, *p* = 0.0133) but, gephyrin-VAT co-localization (t test, *t*_(58)_ = −3.6675, *p* = 0.0005) was decreased due to PME exposure (**Figure 4M,N**). For L2/3 and L5, PME decreased the density of functional GABAergic synapses. Together, the net effect of both glutamate and GABA synapses suggests a net strengthening of glutamatergic synapses in M1, across multiple cortical layers.

### Multi-Region Differential Protein and Phosphopeptide Expression

The combination of protein, phosphopeptide, and anatomical alterations in synapse structure and function in M1 provoked us to ask if there were other brain regions that displayed similar changes due to PME. We updated our analyses of prior proteomic and phosphoproteomic datasets using searches of more up-to-date databases to determine if S1, DLS, and DMS displayed similar changes in protein and phosphopeptide alterations (Grecco, Huang et al. 2022, Grecco, Munoz et al. 2022). We preformed the identical quantitative proteomic and phosphoproteomic analyses that we did in M1. In S1, we identified 10,818 proteins and 2,669 phosphopeptides in our samples, but only 1,637 proteins and 263 phosphopeptides were differentially abundant (**Figure 5A,B**). Similar to the M1 proteome and phosphopeptides, S1 showed a large number of significantly down-regulated proteins compared to up-regulated proteins, and an even balance of up-regulated and down-regulated phosphopeptides, which may be most strongly driven by PME males (**Figure 5C,D**). In DLS, we identified 8,588 proteins and 4,606 phosphopeptides in our samples, but only 839 proteins and 381 phosphopeptides were differentially abundant (**Figure 5E,F**). The DLS proteome and phosphopeptides were similarly up- and down-regulated with the largest effects driven by PME, not sex (**Figure 5G,H**). Finally, in DMS we identified 9,411proteins and 4,594 phosphopeptides in our samples, but only 1,304 proteins and 282 phosphopeptides were differentially abundant (**Figure 5I,J**). The DMS proteome, like M1, saw many proteins down-regulated (∼80%) and an even distribution in phosphopeptide changes with PME males being most dissimilar from PSE males for the proteome (**Figure 5K,L)**. Altogether, M1 showed similar patterns of changes with S1 and DMS in that there was a bias towards decreased protein expression, but a more balanced change in phosphopeptides, while DLS showed a more even distribution of both proteins and phosphopeptides. In regard to the total number of individual proteins that had their abundance altered by PME, S1 showed the largest PME-induced effect. DMS and M1 were nearly equivalently affected and DLS had the least changes due to PME.

**Figure 5:**
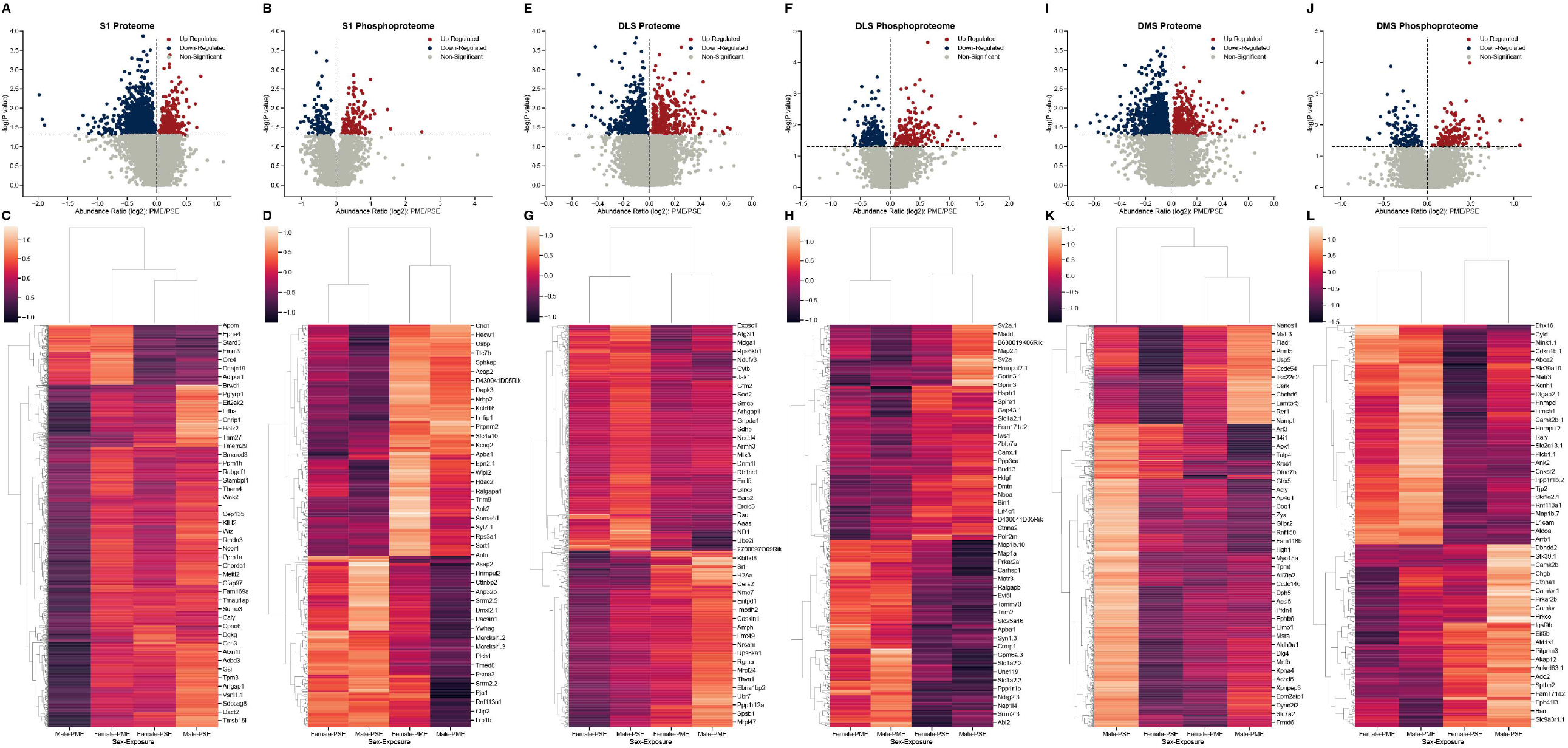
Differential protein and phosphopeptide expression in the primary somatosensory cortex (S1), dorsolateral striatum (DLS), and dorsomedial striatum (DMS) of prenatal methadone exposed offspring. Volcano plots for the differential S1 proteome (A) and phosphoproteome (B) with blue circles representing individual proteins/phosphopeptides decreased in PME vs. PSE and red circles representing individual proteins/phosphopeptides increased in PME vs. PSE which reach the level of significance. Clusterplots of differentially abundant S1 proteins (C) and phosphopeptides (D) with example GeneIDs. Volcano plots for the differential DLS proteome (E) and phosphoproteome (F) with blue circles representing individual proteins/phosphopeptides decreased in PME vs. PSE and red circles representing individual proteins/phosphopeptides increased in PME vs. PSE which reach the level of significance. Clusterplots of differentially abundant DLS proteins (G) and phosphopeptides (H) with example GeneIDs. Volcano plots for the differential DMS proteome (I) and phosphoproteome (J) with blue circles representing individual proteins/phosphopeptides decreased in PME vs. PSE and red circles representing individual proteins/phosphopeptides increased in PME vs. PSE which reach the level of significance. Clusterplots of differentially abundant DMS proteins (K) and phosphopeptides (L) with example GeneIDs. (n = 8 (4M:4F) PME, 8 PSE (4M:4F)).

### Multi-Region Gene and Gene Ontology Functional Enrichments

We also sought to understand the overlap between the PME-altered proteome and phosphoproteome in other brain regions. Comparing the M1 overlap, which implicated significant changes in synapse structure and activity, to the overlap in these other brain regions would allow us to understand the uniqueness of PME-induced protein and phosphopeptide changes in each region in the context of altered associated genes, and alterations in biological processes. Similarly, we performed gene and gene ontology analyses on S1, DLS, and DMS regions for both the proteome and phosphoproteome. For the proteome of all brain regions, only one associated gene target was shared (**Figure 6A**), Pdxk which encodes pyridoxal kinase that converts inactive B6 vitamers into the active cofactor (Keller, Mendoza-Ferreira et al. 2020). Further, the overlap of shared GeneIDs between the proteomes M1-SI (2.76%), M1-DMS (2.51%), M1-DLS (3.47%) were extremely low, suggesting that PME alters the proteome of different brain regions in unique ways (**Figure 6A**). For the phosphoproteome of all brain regions there was a larger overlap of 19 associated GeneIDs (**Figure 6B**). The GeneIDs of the top 5 most positively and negatively altered log_2_(abundance ratios) in the phosphoproteome returned items such as Ank2 and Cldn11, which were also shown in the M1 overlap and found to be altered globally (**Figure 6B**). Also, there were additional associated GeneIDs not found the in M1 overlap that were found in the global phosphoproteome overlap, such as a Add2, Map2, Pclo, Rims1, Bsn, Cacnb4, and Camk2b which are known to alter synapse function and maintenance (Caddick, Wang et al. 1999, Fenster and Garner 2002, Mittelstaedt, Alvarez-Baron et al. 2010, Lisman, Yasuda et al. 2012, Waites, Leal-Ortiz et al. 2013, Kandimalla, Manczak et al. 2018). These data suggest that PME similarly alters the phosphoproteome globally, but the lack of these GeneIDs being returned in the global proteome suggests that M1, and other brain regions, have unique susceptibility to PME.

**Figure 6:**
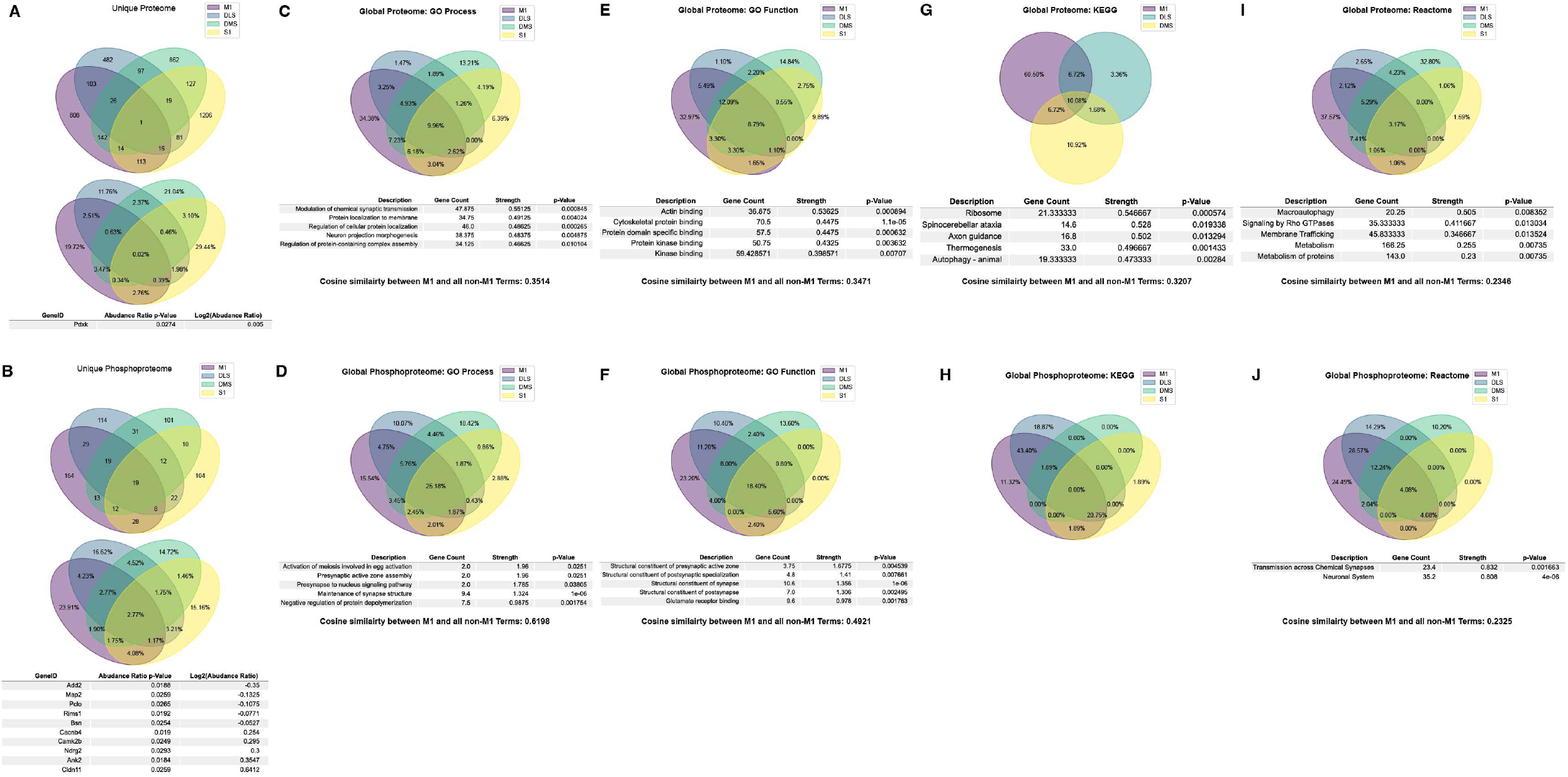
Overlap in the protein, phosphopeptides, and gene ontology enrichments between prenatal methadone exposed primary motor cortex (M1), primary somatosensory cortex (S1), dorsolateral striatum (DLS), and dorsomedial striatum (DMS). Across the global proteome (A), only 1 associated GeneID was conserved across all brain regions, and across the global phosphoproteome (B), 19 associated GeneIDs were altered, with the top 5 strongest positive and negative log2(abundance ratio) GeneIDs listed. For the GO Process gene ontology enrichments, 9.96% of proteins (C) were similar across all brain regions and M1 to non M1-region terms were assessed using cosine similarity and were 35.14% similar. For the phosphoproteome (D), 25.18% of terms were similar and the M1 to non-M1 similarity was 61.98%. For GO Function enrichments, 8.79% of protein terms (E) were similar with a M1 to non-M1 similarity score of 34.71%, and 18.40% of phosphopeptide terms (F) were similar with a M1 to non-Ma score of 49.21%. For KEGG enrichments, there were not any significantly enriched proteins (G) in the S1 region, so the similarity between protein terms was 10.08% and the M1 to non-M1 similarity score was 32.07%. The KEGG enrichments for the phosphoproteome (H) had no overlapping terms and therefore no M1 to non-M1 similarity score. For the Reactome enrichments for the proteome (I) there were 3.17% similarity between protein terms, and a M1 to non-M1 similarity score of 23.46%. For the phosphoprotein Reactome (J) there was 4.08% overlap and a M1 to non-M1 similarity score of 23.25%.

We also looked at gene ontology term overlaps between all brain regions and compared M1 terms to the overlap of all other terms in attempt to quantitatively score how similar changes in M1 were to other regions of interest. For the global proteome, GO Processes were 9.96% similar, with the top 5 strongest terms focusing on synaptic transmission, protein localization, and neuronal assemblies (**Figure 6C**). The GO Process terms for M1 were also compared to the set of all non-M1 brain region terms. This list of terms was vectorized and the cosine similarity between the two vectors were computed, which resulted in a similarity of 0.3415, which can be conceptualized as 34.15% similar for GO Processes for the M1 proteome compared to all other brain region proteomes (**Figure 6C**). For the global phosphoproteome, GO Processes were 25.18% similar, with the strongest relevant terms involving synapse structure and activity (**Figure 6D**). M1 to non-M1 similarity was also very high, 61.98%, which mirrored the differences seen between the global proteome and global phosphoproteome for associated GeneIDs (**Figure 6D**). For GO Function terms in the global proteome, there was an overlap of 8.79% with the strongest terms generally referring to protein binding, with a M1 to non-M1 similarity of 34.71% (**Figure 6E**). In the global phosphoproteome, GO Function was 18.4% similar with terms strongly associated with synapse structure, and glutamate receptor binding, and M1 to non-M1 similarity was 49.21% (**Figure 6F**). For KEGG terms in the global proteome, there was no significant S1 KEGG enrichments to report, so the global overlap contains M1, DLS, and DMS only. There was a 10.08% overlap between these terms, with axon guidance being the relevant strongly associated term (**Figure 6G**). For the KEGG global phosphoproteome, there was no overlap between terms, therefore there was also no M1 to non-M1 similarity score, which can be interpreted as zero (**Figure 6H**). Finally, the global protein Reactome had 3.17% similarity with protein metabolism and membrane tracking being the most relevant strongly enriched terms, with a M1 to non-M1 similarity score of 23.46% (**Figure 6I**). The global phosphoproteome had 4.08% similarity with synaptic transmission and the neuronal system being significantly enriched, with a M1 to non-M1 similarity score of 23.25% (**Figure 6J)**. For a full detailed list of differentially abundant proteins and phosphopeptides for each brain region, please Supplementary Files for S1, DLS, and DMS regions.

## Discussion

Together these data suggest that PME-induced changes to the M1 proteome are uniquely different from the S1, DLS, and DMS proteomes. The average cosine similarity between all gene enrichments is 31.35%, and while some significantly enriched terms generally point to differences in neuronal excitability and protein tracking, only M1 terms return narrow, specific enrichments for glutamate synapses. Also, our protein expression assessments identified changes in specific glutamate receptors, such as ionotropic AMPA and NMDA glutamate receptor subunits (GluA2, GluA3, GluA4, GluN1, GluN2a, and GluN2b) as well as metabotropic glutamate receptors (mGluR1, mGluR2, mGluR3, mGluR5, and mGluR8). Consistent with glutamate signaling proteins being especially affected in M1, we also saw an increase in functional glutamate synapse density in M1, which we did not find in our prior study of S1 (Grecco, Huang et al. 2022). We previously found enhanced glutamate input from layer2/3 M1 neurons to layer 5 M1 pyramidal neurons (Grecco, Mork et al. 2021). The enhanced glutamate synapse density aligns with this. However, the predominant effect of PME on glutamate receptor and glutamate synapse-related proteins was a decrease in expression. Subcellular localization and further functional analyses are needed to determine the interplay of glutamate protein expression and altered synaptic function. Of particular interest, the most differentially expressed protein in M1 as a result of PME is the alpha2-adrenergic receptor, a target of clonidine, which is an adjunct medication used to treat NOWS (Merhar, Ounpraseuth et al. 2021). This receptor also regulates HCN channels which mediate the voltage sag we identified as being enhanced by PME in M1 pyramidal neurons (Sheets, Suter et al. 2011, Grecco, Mork et al. 2021). Relatedly, PME decreased HCN1 channel expression as well as filamin A and Trip8b, two proteins that regulate HCN expression and function (Peters, Singh et al. 2022). One of the functions of HCN channels is to filter glutamate transmission, which may also contribute to the altered glutamate transmission we discovered previously (Grecco, Mork et al. 2021). Like our findings of enhanced glutamate transmission, but decreased glutamate receptor expression, the paradox of increased voltage sag, but decreased expression of proteins that mediate that phenomenon will require deeper investigation. Why glutamate signaling-related protein expression in M1 is so profoundly affected by PME relative to other brain regions is a mystery that will require much more study.

We found a decrease in the density of VGAT-gephyrin co-localization in both of the layers of M1 we analyzed, suggesting a decrease in functional GABA synapses. An assessment of the distribution of the data in regard to sex seemed to indicate that this may be driven by males, although future work is needed to more rigorously assess this. Gene ontology and pathway enrichment analyses did not overwhelmingly show that PME uniquely affected GABAergic synaptic proteins like they did for glutamatergic synapses. Nevertheless, we did find some changes in GABAergic synapse-related protein expression. Consistent with the reduction of functional GABAergic synapses, gephyrin expression was decreased by PME as was the vesicular GABA transporter GAT3 and various GABAA and GABAB receptor subunits.

Interestingly, PME effects in the phosphoproteome of all overlapping brain regions and in the M1 phosphoproteome compared to other non-M1 brain regions generalize more broadly. PME similarly affects phosphopeptides across many brain regions by altering the expression of synaptic proteins that mediate neuronal signaling. Further, the average similarity between M1 and non-M1 enrichments is 44.81%. Yet, while greater similarities exist between M1 and other brain regions for the phosphoproteome than do similarities between the M1 proteome and other brain regions, the number (164) and percentage (23.91%) of differentially abundant unique phosphopeptides is greater than all three other brain regions, suggesting that the magnitude in which PME can uniquely alter the phosphoproteome is greatest in M1 than any other brain region sampled (**Figure 6B**). These data suggest that M1 might be a primary site of PME interaction in the brain, and future studies are needed to delve more mechanistically to understand how this occurs. Although, the current analyses may provide starting points for determining which specific protein and phosphopeptides may be susceptible to PME, and how these may serve as a starting point for intervention, both experimentally and translationally.

There are some noteworthy limitations to bear in mind when considering this multi-omics analysis. First, although the differential protein/phosphopeptide expression in PME offspring indicated alterations in synaptic signaling were present, these enrichment analyses do not provide “directionality.” For instance, while the protein/phosphopeptide abundances in PME offspring indicate that “Glutamate Receptor Activity” is enriched, a GO enrichment analysis does not tell us that glutamate activity is increased or decreased, but only that this is significantly represented given the list of differentially abundant proteins/phosphopeptides. Our data regarding decreased glutamate receptor expression, but increased glutamate synapse density and glutamate transmission is a perfect example of the limitations in trying to interpret functional changes from proteomics data.

Additionally, one-to-one comparisons between proteomics/phosphoproteomics data with synaptic marker findings remain difficult as bulk M1 tissue was taken for the multi-omics analysis. Therefore, the quantified proteins/phosphopeptides may have originated in various M1 layers, glia cells, interneurons of M1, or even presynaptic inputs from other brain regions.

In summary, our findings indicate PME induces prominent disruptions in M1 as well as other anatomically and functionally connected brain regions. We also investigated thousands of proteins and phosphopeptides across brain regions that display differential abundance in PME offspring with functional enrichment in several relevant pathways including those related to synaptic transmission and assembly. These findings suggest that deficits in motor behavior function that persist throughout the lifespan due to POE may result from persistent neuroadaptations in M1 and connected brain regions induced by opioid exposure during fetal development. Deciphering how POE produces these changes may lead to novel treatments for mitigating the negative impact of POE on motor and other features of development.

## Data Availability Statement

The raw data supporting the conclusions of this article will be made available by the authors without undue reservation. All supplementary files and raw data can be found at https://github.com/dlhagger/PME-MultiRegion-Proteomics.

## Ethics Statement

The animal study was reviewed and approved by Indiana University School of Medicine Institutional Animal Care and Use Committee.

## Author Contributions

GG, J-YH, ED, AM, HCL, and BA designed experiments. GG, generated animals. ED and AM performed proteomics and phosphoproteomic studies. DLH completed proteomic and phosphoproteomic analyses. J-YH completed all immunostaining work. All authors discussed the results and contributed to all stages of manuscript preparation.

## Figure Legends

**Supplemental Figure 1:**
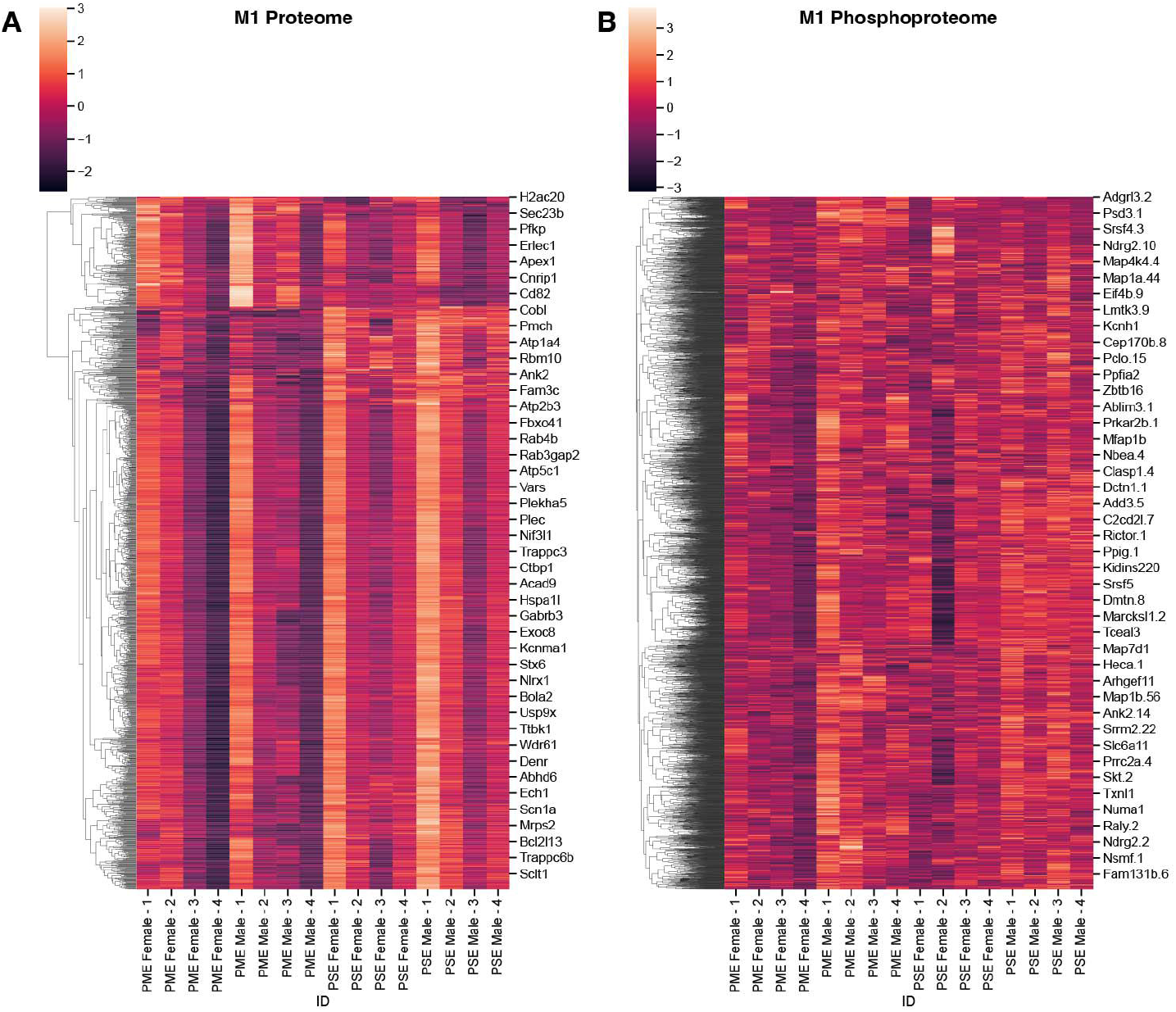
Differential protein and phosphopeptide expression in the primary motor cortex (M1) of prenatal methadone exposed offspring. Clusterplots of differentially abundant proteins (A) and phosphopeptides (B) with example GeneIDs. (n = 8 (4M:4F) PME, 8 PSE (4M:4F)).

